# Characterization of Bacterial Isolates in Vermicompost Produced from a Mixture of Cow dung, Straw, Neem leaf and Vegetable Wastes

**DOI:** 10.1101/2020.07.01.183467

**Authors:** Jyotismita Satpathy, Mirkashim H. Saha, Aditya S. Mishra, Sujit K. Mishra

## Abstract

Vermicomposting is a non-thermophilic biological oxidation process of composting where certain species of earthworm are used to enhance the process of conversion of organic waste to compost. Earthworm helps in influencing the growth of certain microbial species, and also improves the physical and chemical properties of the soil. The microbial population present in vermicompost play an important role in increasing the productivity of crop as well as maintain the structural stability of the soil. Different types of bacteria found in vermicompost and is depends on the environmental condition and the raw materials used in vermicomposting. Owing to this, a study was carried out to identify the bacteria exist in vermicompost produced from cow dungs, straw, neem leafs and vegetable wastes. The phenotypic studies such as colony morphology, microscopic studies, and biochemical characterization have identified eight bacterial species namely *Actinomyces israelli, Azotobacter, Micrococus luteus, Bacillus cereus, Bacillus subtillis, Pseudomonas aeruginosa, Enterobacter* in the vermicompost. All these bacteria were present in the gut of *Eesenia Foetida* and found beneficial for the soil and crop plants.

## INTRODUCTION

Microbes are the essence of biodiversity and also play important role in functioning of the ecosystem. Microbes such as bacteria, fungi, actinomycetes, etc. responsible for biochemical degration of organic matter (Emperor et al., 2015) and maintains ecological balance. Vermicompost are the stabilize and non thermophilic products, that are produced by interactions of earthworms and microorganisms that rich in microbial activity. Vermicomposting is one of the easiest methods to recycle agricultural wastes into product like good quality of compost. The compost is rich in nutrients, growth promoting substances, and beneficial microbes. It is a major component of organic farming system. Earthworm is the crucial driver of vermicompost that aerate, alter the microbial activity and their degradation potential (Fracchia et al., 2006). Vermicompost improves the soil aggregation, soil fertility, plant nutrition and also growth of beneficial microbes (Pereira et al., 2014). It improves soil aeration and water holding capacity. It suppresses diseases in plant and help to enhances plant growth, restores microbial population which includes nitrogen fixers, phosphate solubilizers, etc. and also provides macro and micro nutrients to the crop plants. It also helps to improve structural stability of the soil which helps prevent soil erosion (Zhu et al., 2017) and ultimately increases productivity of different crops (Khan et al., 2011).

In addition, vermicomposting also improves the physiochemical and biological properties of soils and enhances the diversity and population of beneficial microbial communities (Pathma et al., 2012). Earthworms ingest soil microbes with organic residues from soil and during passage through the intestinal tract, their population may increase. The ingested microbes along with wastes enter the gut and play an important role in earthworm nutrition by donating microbial enzymes that help in breakdown of organic matter (Selvi et al., 2015). Thus, identifying the bacterial population in vermicompost could provide insight into the quality of vermicompost produced and knowing the beneficial bacteria involves in the process o composting. Therefore, the present study was carried out with an objective to identify the bacterial species involves in the vermicomposting of organic and agricultural wastes such as cow dung, straw, neem leaf and vegetable wastes.

## MATERIALS AND METHODOLOGY

### Preparation of vermicompost

Pit method of vermicomposting was followed where composting is done in the cemented tanks. The vermicomposting tank was established in a cool and moist place, and a shade was was established over the tank. Waste vegetable peels, neem leaves, dried straw or grass were sprinkled with cowdung slurry for quick decomposition. The materials which were under the sun for about 7-10 days or partially decomposed were placed in the tank in alternative layers respectively. Requisite numbers of red earthworms (*Eisenia Foetida*) were released into the tank over the mixture. The compost mixture in the tank was covered with gunny bags. Water sprinkling was done regularly as per requirement and 70-80 % moisture was maintained.

### Bacterial sample preparation

Two gram of vemicompost was collected from the vermicomposting unit located at CUTM campus, Balangir, Odisha, India in a sterilized beaker and transported to the laboratory for isolating and identifying the bacterial species in the vermicompost. One gram was taken separately in a sterilized conical flask containing 100 ml of sterile distilled water and mixed well. One ml of diluted sample was taken and poured into 9 ml of sterile distilled water and then the sample serially diluted (Selvi et al., 2015) for culturing.

### Isolation of bacteria from vermicompost

For isolating bacteria from vermicompost, nutrient agar media was used that was consists of Agar Agar (type-I) and Nutrient broth. Pour plate method was followed for isolation in which the serial diluted samples were poured on respective petriplates containing the nutrient agar media and then incubated at 37 °C for 24 hours in a bacteriological incubator (Selvi et al., 2015). After incubation the appeared bacterial colonies were subjected to subculturing for characterization.

### Characterization and identification of bacteria

The bacterial strains isolated were characterized on the basis of their morphological, microscopic and biochemical properties. The colony morphologies were then studied on naked eye. The parameters taken were colour of the colony, solubility on water, opacity, elevation, texture and consistency. After that, microscopic study was done by gram staining method and then visualized under microscope. Further, the bacterial strains were identified by performing biochemical test that included IMViC tests (Indole, Methyl red, Voge’s Proskauer, Citrate), Oxidase test and TSI test (Triple sugar iron) (Olutiola et al., 2000). The reagents used for the biotyping were obtained from Hi-Media.

## RESULTS

### Characterization of the isolated bacteria

Different type of bacterial colonies appears in the agar culture plates. The colonies were found with different colour, size, texture, elevation, edge, and shape, etc. The isolated bacterial strain were characterized by colony morphology study, microscopy study and biochemical characterization.

### Colony Morphology Study

There were eight different types of colonies observed in primary culture subjected for sub-culturing by streaking methods. The colonies were coded as BIFV A, BIFV B, BIFV C, BIFV D, BIFV E, BIFV F, BIFV G, BIFV H. The result of streak plates on the culture plates were depicted in Figure 1. The colony of BIFV A was white in colour, opaque, umbonate with rough in texture (Slack et al., 1969). The colonies of BIFV B were white in colour and were opaque, rised and smooth in texture having irregular shape. BIFV C showed yellow colour colony which was circular in shape having lobulate edge. The colony of BIFV D was white with curled edge and rough texture. BIFV E was white colony having rised elevation and lobulate edge. The BIFV F formed white colonies having filamentous edges and flat elevations. BIFV G was white colour colony having circular shape with convex elevation. The colony of BIFV H was white in colour having mucus and entire edge with circular shape. The characteristic morphology are present below in Table 1.

**Table 1.** The colony morphology of isolated bacterial strains.

| Code | Colour | Opacity | Elevation | Texture | Consistency | Shape | Edge |
| --- | --- | --- | --- | --- | --- | --- | --- |
| BIFV A | White | Opaque | Umbonate | Smooth | Stick to loop | Irregular | Undulate |
| BIFV B | White | Opaque | Rised | Smooth | Stick to loop | Irregular | Lobulate |
| BIFV C | Yellow | Opaque | Umbonate | Rough | Stick to loop | Circular | Undulate |
| BIFV D | White | Opaque | Umbonate | Rough | Stick to loop | Irregular | Curled |
| BIFV E | White | Opaque | Rised | Smooth | Stick to loop | Irregular | Lobulate |
| BIFV F | White | Opaque | Flat | Rough | Sticky | Irregular | Filamentous |
| BIFV G | White | Opaque | Convex | Smooth | Sticky | Circular | Entire |
| BIFV H | White | Opaque | Flat | Smooth | Mucus | Circular | Entire |

**Figure 1.**
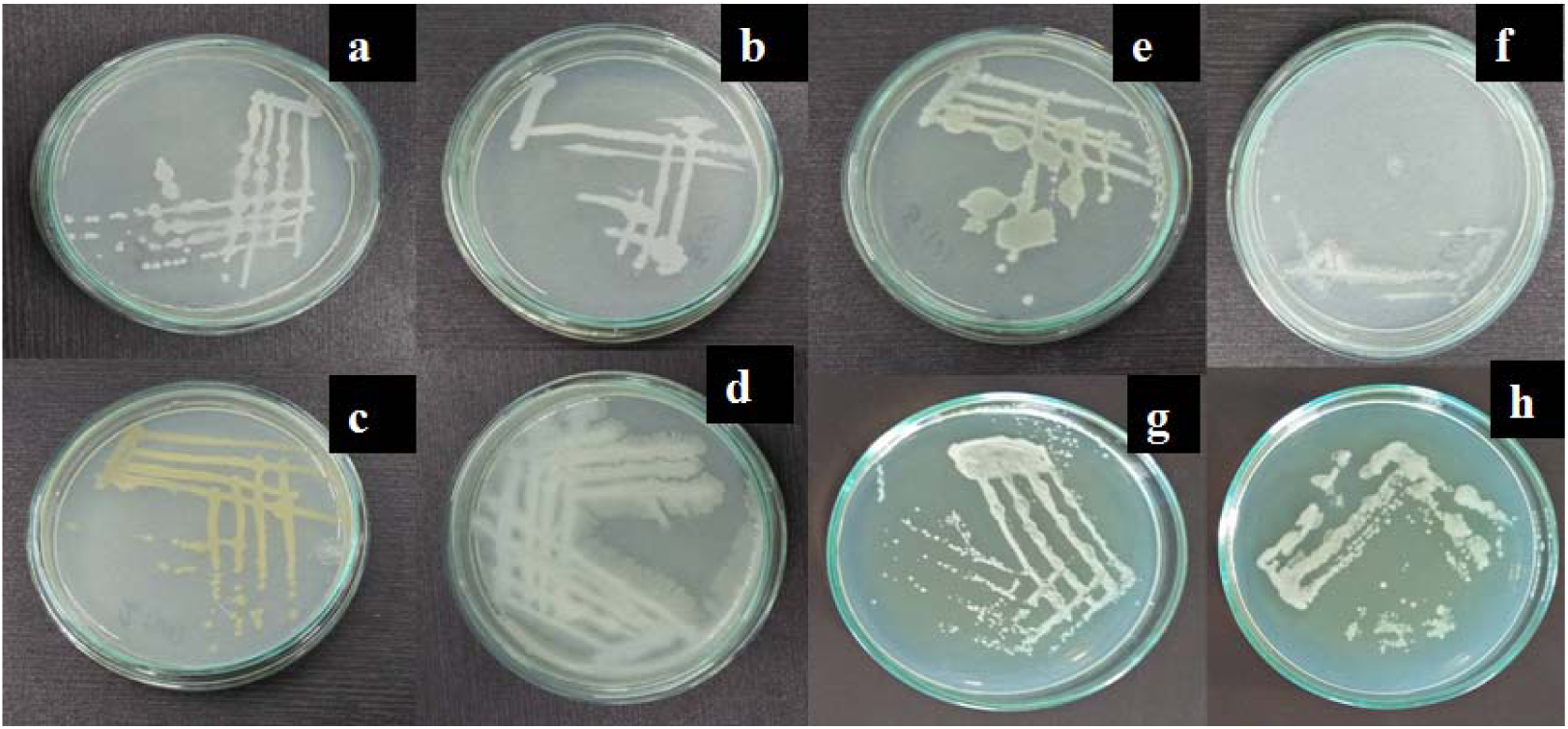
Depicting the streak plates. (a) BIFV A (White), (b) BIFV B (White), (c) BIFV C (yellow), (d) BIFV D (White), (e) BIFV E (White), (f) BIFV F(White), (g) BIFV G (White), (h) BIFV H (White)

### Microscopic Study

The bacterial strains isolated from vermicompost sample were visualized under the compound microscope and the results are represented in Table 2. The BIFV A, BIFV B, BIFV D, BIFV E, BIFV F were observed as Gram’s positive bacterial strain with coccus form. The BIFV C was Gram’s positive bacterial strain with rod shape. BIFV G was Gram’s negative bacterial strain with coccus shape but BIFV H was also Gram’s negative strain with rod shape. Hence, the result obtained from microscopic study indicates about occurrence of Gram’s positive as well as Gram’s negative bacterial strains.

**Table 2.** Responses of isolated bacterial strains to Gram’s stain.

| Sl. No. | Code | Gram's Response | Shape |
| --- | --- | --- | --- |
| 1. | BIFV A | +ve | Coccus |
| 2. | BIFV B | +ve | Coccus |
| 3. | BIFV C | +ve | Rod |
| 4. | BIFV D | +ve | Coccus |
| 5. | BIFV E | +ve | Coccus |
| 6. | BIFV F | +ve | Coccus |
| 7. | BIFV G | -ve | Coccus |
| 8. | BIFV H | -ve | Rod |

### Biochemical characterization

The biochemical tests results were interpreted by the reaction occur in the form of change in colour that was observed in naked eye and shown in Figure 2. In Indole test, the strain of BIFV A, BIFV B, BIFV C, BIFV D,BIFV E, BIFV F, BIFV G, BIFV H showing pale yellow colour or slightly cloudy that indicates the Indole was not produced from tryptone that shown negative result (MacWilliams et al., 2012). As Methyl red test was used to detect the fermentation of glucose which indicated by the formation of red colour (Hemraj et al., 2013). Here, BIFV A, BIFV B, BIFV C, BIFV E were shown positive result and BIFV C, BIFV F, BIFV G, BIFV H were shown negative results. BIFV A, BIFV C, BIFV D, BIFV H shown positive result for VP test by producing pink colour but BIFV B, BIFV E, BIFV F, BIFV G were negative. In case of Citrate test, BIFV A was shown negative result whereas BIFV B, BIFV C, BIFV D, BIFV E, BIFV F, BIFV G, BIFV H were producing the blue colour which referred to positive result. In Oxidase test, the bacterial strain BIFV A, BIFV B, BIFV C, BIFV E, BIFV F, BIFV G, BIFV H shown positive result whereas BIFV D shown negative for oxidase test. In the Triple Sugar Iron (TSI) test, the bacterial strain namely BIFV A,BIFV B, BIFV C, BIFV D, BIFV E, BIFV F, BIFV G were detected as the fermenters of glucose only but BIFV H were detected as glucose, lactose and sucrose by producing yellow colour butt and slant which indicates acid production. The results were depicted in Table 3.

**Table 3.**
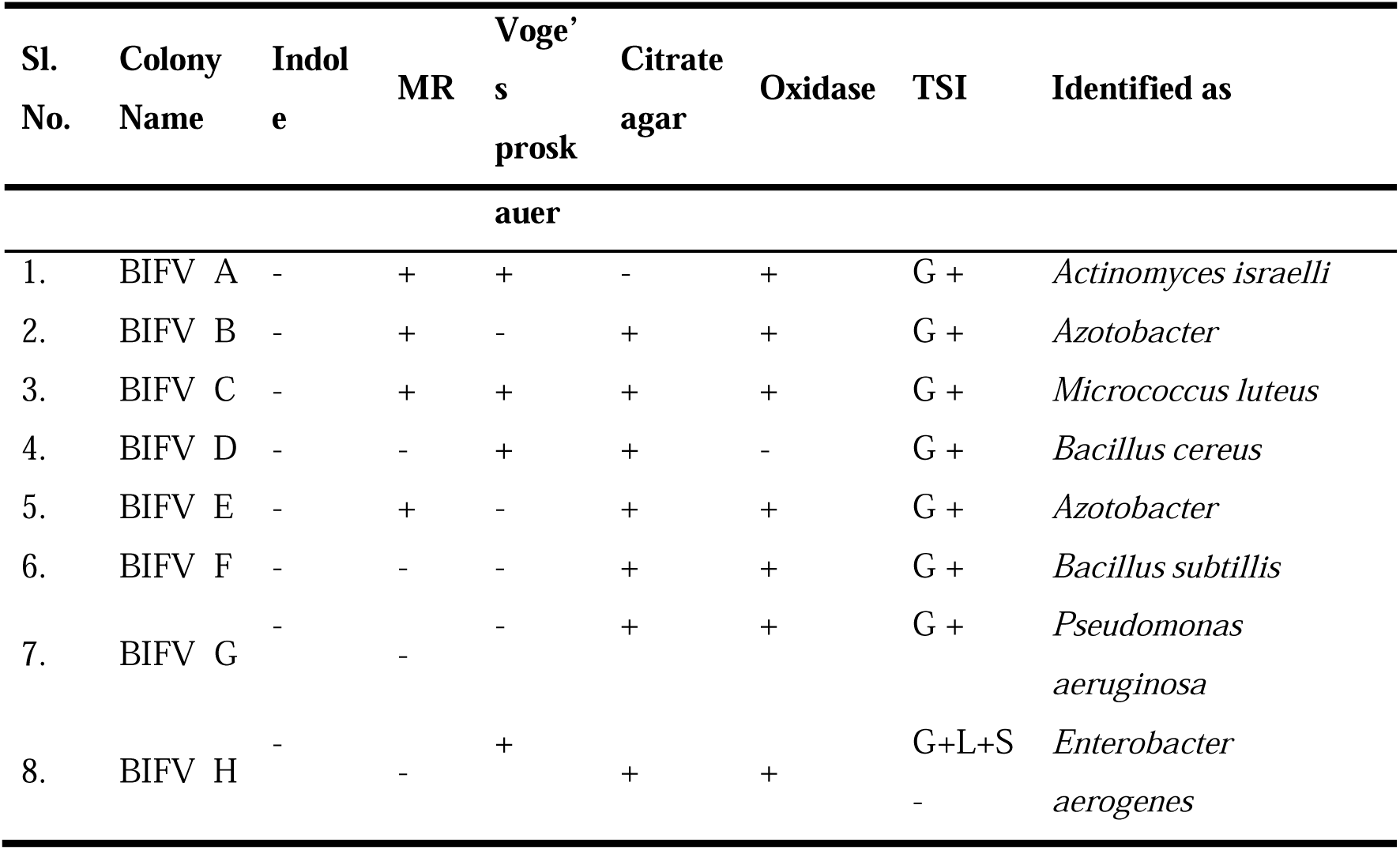
Results of Biochemical test carried out for isolated bacterial strains.

**Figure 2.**
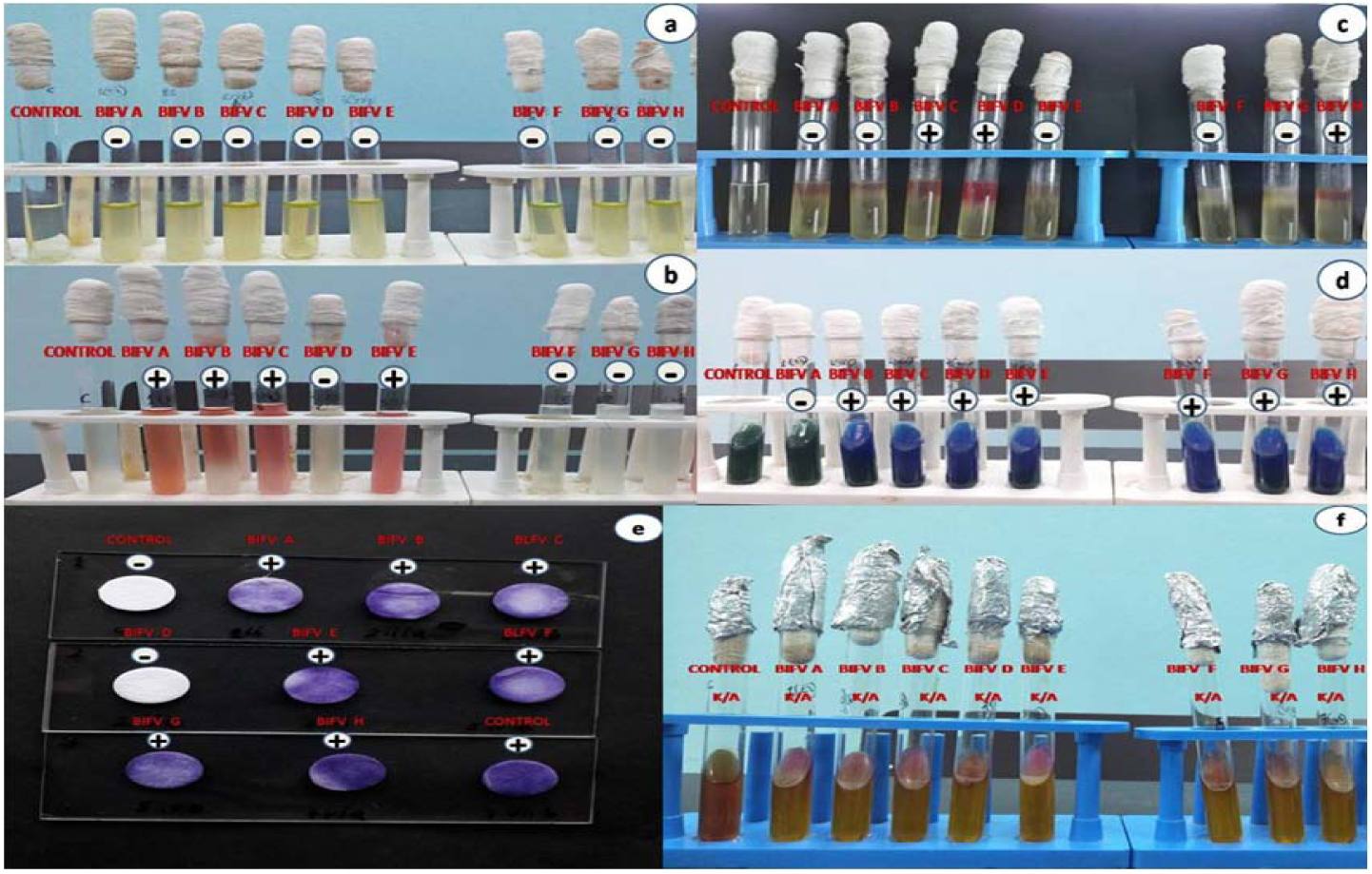
Biocemical tests results. (a) Results of Indole test, Control tube is on extreme left. Negative result is indicated by formation of pale yellow or slightly cloudy ring coloured ring; (b) Results of Methyl Red test. Control tube is on extreme left. Positive result indicates formation of red colour; (c) Result of Voge’s Proskauer test. Control tube is on extreme left. Positive result is indicated by formation of pink colour; (d) Result of Citrate Test. Control tube is on extreme left. Positive result is indicated by formation of blue colour; (e) Results of oxidase test. Positive Control is on extreme right. Positive result is indicated by formation of dark blue colour; (f) Results of TSI test. Positive Control is on extreme left. K/A indicates Alkaline slat and Acidic butt. And A/A indicates Acidic slant and Acidic butt.

## DISCUSSION

Isolation of bacterial strains from vermicompost helps to identify the microbes that enhancing soil fertility and also help in crop improvement. By identifying bacteria we know their role in the plant growth and development, and promote the growth of those bacteria which have beneficial role in vermicompost to enhance the nutrient quality in vermicompost. On the basis of morphological, microscopic and biochemical studies, eight bacterial strains were primarily identified viz. *Actinomyces israelli, Azotobacter, M. Luteus, B. Cereus, B. Subtillis, P. aeruginosa* and *Enterobacter*.

The BIFV A was supposed to be a strain of *Actinomyces israelli* as the colour of the colonies was colourless or white having umbonate elevation with irregular, opaque in opacity. This was congruent with the result obtained by Slack et al., 1969. The BIFV B and BIFV E were supposed as *Azotobacter* as these produced white colonies having irregular shape. The result was congruent to result obtained by Upadhyay et al., 2015. The BIFV C was predicted as *Micrococcus luteus* due to production of yellow colonies according to the result obtained by Wieser et al, 2002. The BIFV D was proposed as *Bacillus cereus* as it was having irregular shape with lobulate edge. The BIFV F was proposed as *Bacillus subtillis* having sticky in nature. This result was same as result of Yunping et al., 2019. The BIFV G was proposed as stain of *Pseudomonas aeruginosa* as having convex elevation, opaque, and circular shape, and same as result of Kirists et al., 2005. The BIFV H was proposed as *Enterobacter arrogenes* strain as having mucus consistency with colourless or white colony (Stiles et al., 1981).

The results obtained from microscope study indicated both Gram positive and Gram negative bacterial strain with coccus and bacillus forms present in vermicompost. Selvi et al. (2015) also reported the same in vermicompost sample. However, Begum et al. (2018) were reported that the bacterial strains were gram positive in nature and rod in shape.

The biochemical test results were interpreted by the reaction that taken place in form of the change in colour (Figure 2). In the Indole test, the strain BIFV A, BIFV B, BIFV C, BIFV D, BIFV E, BIFV F, BIFV G and BIFV H were unable to produce indole from tryptone both giving negative result. As methyl red test used to detects fermentation of glucose that indicated by the production of red colour. Here the BIFV A, BIFV B, BIFV C, BIFV E were detected as good fermenter of glucose where as BIFV D, BIFV F, BIFV G and BIFV H were glucose non fermenter. In VP test, BIFV A, BIFV C, BIFV D, BIFV H were showing positive result but BIFV B, BIFV E, BIFV F, BIFV G were showing negative result. In case of citrate agar test, BIFV B, BIFV C, BIFV D, BIFV E, BIFV F, BIFV G were producing blue colour which showed positive result and utilization of citrate as the sole source of carbon (Van Hofwegen et al., 2016). But BIFV A did not change colour so it shows negative result. In oxidase test, the BIFV A, BIFV B, BIFV C, BIFV E, BIFV F, BIFV G, BIFV H were producing blue to purple colour showing positive result. But the BIFV D was shown negative result. The oxidase activity should be performed to provide the increased accuracy in characterization and differentiation of member of *Micrococcaceae* family (Baker, 1984). The TSI identified BIFV A, BIFV B, BIFV C, BIFV E, BIFV F, BIFV G as fermenter of glucose only which is indicated by production of yellow colour butt and red slant which was pointed out towards alkaline production. But BIFV H was fermenter of glucose, lactose and sucrose which indicated by production of yellow butt and slant which pointed out acid production.

## CONCLUSION

Vermicomposting is a bio-oxidative process in which the earthworm interacts intensively with the microorganism that accelerating the stabilization of organic matter, and resulting in improved growth and yield of crop plants. It is cost-effect and ecofriendly waste management technology that improves physiochemical and biological properties of agricultural soil. The study supports the presence of a group of bacterial strains in the vermicompost produced from vegetable wastes, cowdung, neem leaf extracts and straw. The bacterial strains identified by phenotypic studies such as colony morphology, microscopic study and biochemical characterization were *Actinomyces israelli, Azotobacter, Micrococcus luteus, Bacillus cereus, Azotobacter, Bacillus subtillis, Pseudomonas aeruginosa*, and *Enterobacter aerogenes*. Further characterization with 16S rDNA sequencing is needed for confirmation and validation of these bacterial strains. This finding states that all these bacteria are beneficial as they enhance the nutrient status of vermicompost as well as improve the soil aeration and fertility.

## ACKNOWLEDGEMENTS

The authors acknowledge the Centurion University of Technology and Management, Odisha, India for the facilities provided to carry out this work and thankful to Mr. Pradeep Sarangi (Regional Director, CUTM, Bolangir) for giving motivation to conduct the study.

